# Pollen feeding proteomics: salivary proteins of the passion flower butterfly, *Heliconius melpomene*

**DOI:** 10.1101/013128

**Authors:** Desiree Harpel, Darron A. Cullen, Swidbert Ott, Chris D. Jiggins, James R. Walters

## Abstract

While most adult Lepidoptera use flower nectar as their primary food source, butterflies in the genus *Heliconius* have evolved the novel ability to acquire amino acids from consuming pollen. *Heliconius* butterflies collect pollen on their proboscis, moisten the pollen with saliva, and use a combination of mechanical disruption and chemical degradation to release free amino acids that are subsequently re-ingested in the saliva. Little is known about the molecular mechanisms of this complex pollen feeding adaptation. Here we report an initial shotgun proteomic analysis of saliva from *Heliconius melpomene*. Results from liquid-chromatography tandem mass-spectrometry confidently identified 31 salivary proteins, most of which contained predicted signal peptides, consistent with extracellular secretion. Further bioinformatic annotation of these salivary proteins indicated the presence of four distinct functional classes: proteolysis (10 proteins), carbohydrate hydrolysis (5), immunity (6), and “housekeeping”(4). Additionally, six proteins could not be functionally annotated beyond containing a predicted signal sequence. The presence of several salivary proteases is consistent with previous demonstrations that *Heliconius* saliva has proteolytic capacity. It is likely these proteins play a key role in generating free amino acids during pollen digestion. The identification of proteins functioning in carbohydrate hydrolysis is consistent with *Heliconius* butterflies consuming nectar, like other lepidopterans, as well as pollen. Immune-related proteins in saliva are also expected, given that ingestion of pathogens is a very likely route to infection. The few “housekeeping” proteins are likely not true salivary proteins and reflect a modest level of contamination that occurred during saliva collection. Among the unannotated proteins were two sets of paralogs, each seemingly the result of a relatively recent tandem duplication. These results offer a first glimpse into the molecular foundation of *Heliconius* pollen feeding and provide a substantial advance towards comprehensively understanding this striking evolutionary novelty.

## 1. Introduction

Most adult Lepidoptera use flower nectar as their primary food source. Nectar is typically rich in water and carbohydrates but quite limited as a source of amino acids(H. G. Baker, 1975; H. G. Baker and I. Baker, 1977; 1973). Consequently, most Lepidopteran species primarily acquire nutritional protein as larvae feeding on leafy plant material, storing nitrogen and essential amino acids for use during pupation and adulthood (Dunlap-Pianka et al., 1977). Intriguingly, a striking exception to this general pattern is found among butterflies in the genus *Heliconius*, the passion flower butterflies. In addition to nectar feeding, adult *Heliconius* butterflies feed on pollen, a trait with a single origin in this genus (Beltran et al., 2007; Brown, 1981; Gilbert, 1972). Pollen has high nitrogen and essential amino acid content, providing *Heliconius* butterflies with a substantial source of nutritional resources typically thought to constrain adult lepidopteran reproduction and longevity(Dunlap-Pianka et al., 1977; Gilbert, 1972; O'Brien et al., 2003). Accordingly, *Heliconius* butterflies are unusually long-lived, with adult life-spans known to last beyond six months (Gilbert, 1972). Females lay eggs at a moderate and continuous rate throughout adulthood without the reproductive or ovarian senescence characteristic of related butterflies. Carbon isotope analysis has demonstrated that essential amino acids from pollen are directly incorporated into eggs, and excluding pollen from adult *Heliconius* results in dramatic reductions of life-span and fecundity (Dunlap-Pianka et al., 1977; O'Brien et al., 2003). Thus pollen feeding clearly represents a remarkable evolutionary innovation that catalyzed dramatic changes in the physiology and life-history of *Heliconius* butterflies. However, many aspects of this adaptation remain enigmatic and in particular it remains unclear how amino acids are captured from the pollen.

*Heliconius* butterflies do not directly ingest pollen grains. Rather, pollen is collected and stored on the outside of the proboscis (Fig. 1), which has an array of unusually dense and long sensory bristles which presumably facilitate pollen collection and retention (Krenn and Penz, 1998). A suite of behavioral adaptations are also associated with pollen feeding, including sophisticated flower handling and a stereotypical coiling-uncoiling of the proboscis that agitates the collected pollen load (Krenn, 2008; Krenn et al., 2009; Penz and Krenn, 2000). During this pollen processing, saliva is exuded from the proboscis into the pollen and ingested some time later, presumably transporting free amino acids back into the butterfly’s digestive tract.

**Figure 1.**
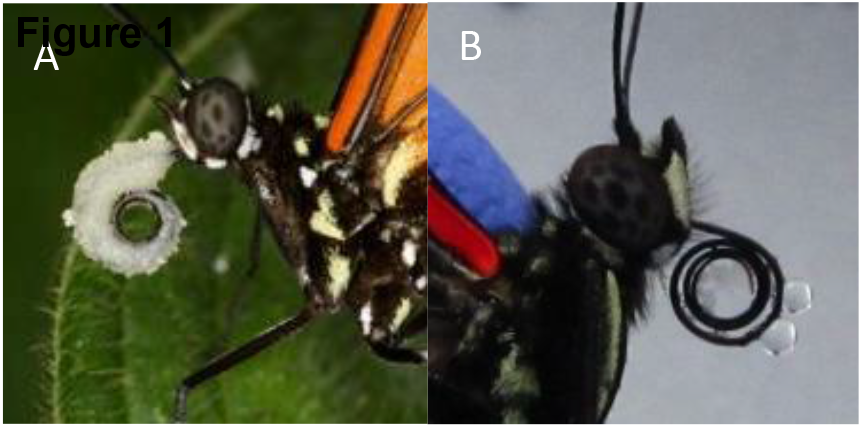
A) Heliconius butterfly with a large load of pollen on the proboscis. B) Saliva droplets exuded onto the proboscis after stimulation with microscopic glass beads during saliva collection.

There has been considerable uncertainty regarding the exact mechanism by which amino acids are released from the pollen grains. Early hypotheses favored a “passive” process. In the initial description of *Heliconius* pollen feeding, Gilbert (1972) suggested that germination of pollen when moistened on the proboscis was sufficient to release free amino acids (Gilbert, 1972). Later Erhardt & Baker (1990)proposed a diffusion process. However, a recent demonstration that proboscis coiling-uncoiling causes substantial mechanical disruption of pollen grains undermines these “passive” hypotheses, indicating instead that *Heliconius* butterflies actively degrade their pollen. Additionally, colorimetric assays of proteolytic activity clearly show *Heliconius* saliva contains proteases that likely degrade pollen enzymatically to complement mechanical disruption (Eberhard et al., 2007). Thus the behavior of pollen processing in saliva acts as an extra-oral digestion (Krenn et al., 2009), but the proteins involved in this process remain unknown.

Here we report an initial investigation into the molecular components of pollen feeding. Using liquid chromatography mass spectrometry (LC-MS) “shotgun” proteomics, we analyzed the protein content of saliva from *Heliconius melpomene*. We confidently identified more than thirty proteins from *Heliconius* saliva, including several putatively secreted proteins with predicted proteolytic function. Also prevalent were proteins predicted to function in carbohydrate hydrolysis and immunity. These results lay the foundation for future investigations into the molecular origins and mechanisms of *Heliconius* pollen feeding.

## 2. Methods

### 2.1 Butterfly care, saliva collection and preparation

*Heliconius melpomene aglaope* were purchased as pupae from commercial providers (Stratford Butterfly Farms, Stratford-Upon-Avon, Warwickshire, UK) and reared in a temperature and humidity controlled greenhouse at the University of Cambridge’s Madingley Field Station, Madingley, UK. Butterflies were kept in cages 1.5 m tall, 1.5 m wide, by 1m deep and provisioned with artificial nectar consisting of 10% sucrose solution in water augmented with 5 g/L Critical Care Formula (Vetark Professional, Winchester UK). In order to minimize contamination of saliva samples with food or pollen proteins, the butterflies were not provided with plants or another pollen source. Additionally, for at least 36 hours before sampling, the Critical Care Formula supplement was removed from the artificial nectar.

Saliva samples were collected by applying a small amount of water-moistened glass beads (<106 μM, Sigma-Aldrich, St Louis, MO, USA) to the proboscis with an insect pin and then washing the proboscis and beads into a 1.5 µL microcentrifuge tube using a pipettor. Typically the application of beads or even just the manipulation of the proboscis with a pin caused visible droplets of saliva to be exuded from the proboscis, usually from the outer edge proximal to the head (Fig 1). The same 150 µL of deionized water was used repeatedly to rinse saliva and beads from the proboscis of 8-10 butterflies per round of collection. Two rounds of collection were performed in one day, separated by 1.5 h, using the same 150 µL diH20. Sampling on two different days provided a pair of biological replicates for proteomic analysis.

Each of the two 150 μL samples was vacuum-centrifuged at 60C to reduce volume to 50 μL. 20 μL per sample was kept for polyacrylamide gel electrophoresis, and the remaining 30 μL was submitted for direct shotgun proteomic analysis via LC-MS.

### 2.2 Protein gel electrophoresis

For polyacrylamide gel electrophoresis, 2.6 vol sample were mixed with 1 vol 4 NuPAGE LDS Sample Buffer (Invitrogen) and 0.4 vol 10 NuPAGE Reducing Agent (0.5 M dithiothreitol; Invitrogen). The samples were heated to 70°C for 10 min, loaded on 4–12% NuPAGE Bis-Tris 1.0mm precast gels (Invitrogen) and electrophoresed in NuPAGE MOPS running buffer at 4 mA/gel for about 100 min. Gels were then fixed and silver stained using standard methods, followed by imaging on a flat-bed scanner.

### 2.3 Mass spectrometry and analysis

Each biological replicate was split into two technical replicates, so a total of four LC-MS experiments were performed. Samples were digested and analyzed *in toto*, one experiment per replicate, without prior gel fractionation. Samples submitted for LC-MS analyses were dried down and resolubilised in 20 mL of 50 mM ammonium bicarbonate. Proteins were then reduced (5 mM DTT) and alkylated (15mM iodoacetamide) before being digested overnight with trypsin. The samples were then dried and resuspended in 20 mL 0.1% formic acid and pipetted into a sample vial and placed in the LC autosampler.

All LC-MS experiments were performed using a nanoAcquity UPLC (Waters Corp., Milford, MA) system and an LTQ Orbitrap Velos hybrid ion trap mass spectrometer (Thermo Scientific, Waltham, MA). Separation of peptides was performed by reverse-phase chromatography using at a flow rate of 300 nL/min and a Waters reverse-phase nano column (BEH C18, 75 mm i.d. x 250 mm, 1.7 mm particle size). Peptides were loaded onto a pre-column (Waters UPLC Trap Symmetry C18, 180 mm i.d x 20mm, 5 mm particle size) from the nanoAcquity sample manager with 0.1% formic acid for 3 minutes at a flow rate of 10 mL/min. After this period, the column valve was switched to allow elution of peptides from the pre-column onto the analytical column. Solvent A was water + 0.1% formic acid and solvent B was acetonitrile + 0.1% formic acid. The linear gradient employed was 5-50% B in 60 minutes.

The LC eluant was sprayed into the mass spectrometer by means of a New Objective nanospray source. All *m/z* values of eluting ions were measured in an Orbitrap Velos mass analyzer, set at a resolution of 30000. Data dependent scans (Top 20) were employed to automatically isolate and generate fragment ions by collision-induced dissociation in the linear ion trap, resulting in the generation of MS/MS spectra. Ions with charge states of 2+ and above were selected for fragmentation. Post-run, the data was processed using Protein Discoverer (version 1.2, ThermoFisher) and converted to mascot generic format (*.mgf*) files for subsequent database searching.

### 2.4 Mass spectra analysis

MS/MS spectra were searched against the *H. melpomene* predicted protein set (downloaded from butterflygenome.org, last updated June 4, 2012) using the Mascot search engine (Perkins et al., 1999). The search parameters were as follows: digestive enzyme-trypsin, maximum missed cleaves-2, fixed modifications-carbamidomethyl, variable modifications-oxidation (M), peptide mass tolerance- 25 ppm, fragment mass tolerance-.8 Da, mass values-monoisotopic, instrument type- ESI-TRAP. The *cRAP* database (via The Global Proteome Machine, www.thegpm.org), last updated February 29, 2012, was also included to search for contaminants in the samples. A false discovery rate (FDR) was calculated by simultaneously searching spectra against a decoy database created by reversing the sequences of the *H.melpomene* protein set. Proteins were identified using peptide and protein identifications validated through Scaffold 4.0 (Searle, 2010). Peptide threshold was established at 90% and protein threshold at 95%, using the Peptide Prophet algorithm and Protein Prophet algorithms respectively, with at least two unique peptide matches required in each sample. Protein and peptide FDR were 0% to ensure high confidence in identifications.

### 2.5 Functional predictions

Proteins identified via LC-MS were functionally annotated bioinformatically using sequence homology. Proteins were searched against the NCBI non-redundant protein database using BLASTP (Altschul et al., 1990). Proteins were also submitted to InterproScan (Zdobnov and Apweiler, 2001). For each protein identified, putative function was manually assigned after reviewing and integrating bioinformatic search results.

## 3. Results and Discussion

### 3.1 SDS-PAGE

Protein electrophoresis revealed a relatively sparse collection of proteins present in the saliva (Fig. 2). Only about 20 distinct bands were visible in the saliva sample. Notably, none of the bands were concordant with bands observed in the dietary supplement, indicating that the saliva was not contaminated with Critical Care Formula diet supplement.

**Figure 2.**
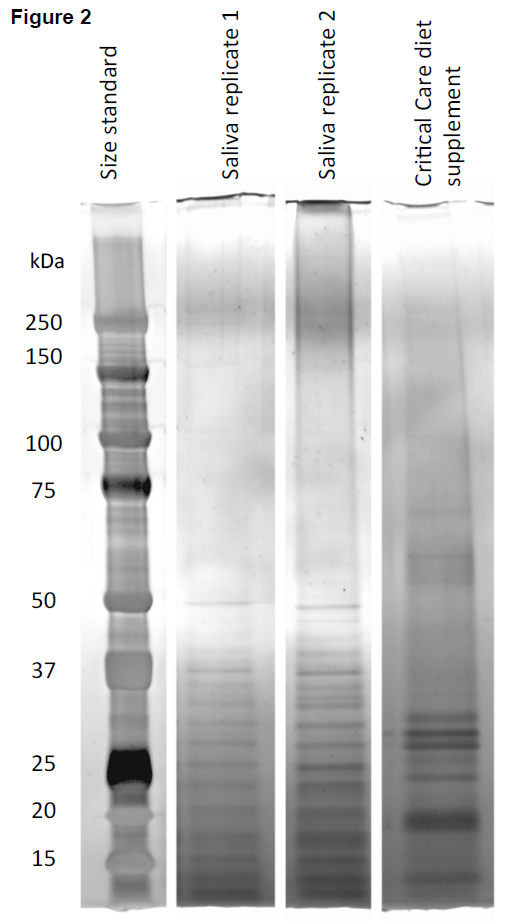
PAGE analysis of *H. melpomene* saliva and Critical Care Formula diet supplement. Size standard is in kiloDaltons (kDa)

### 3.2 Shotgun Proteomics

After filtering the protein hits by significance using Scaffold and removing all contaminant protein hits, a total of 31 proteins were confidently identified from *H.melpomene* adult saliva. Results are summarized in Table 1. There was substantial consistency between biological replicates, with 24 proteins (77%) identified in both samples. Technical replication was also good, with 22 proteins (70%) identified in all four replicates. We also identified and discarded a few obvious contaminant proteins in the filtered LC-MS results (e.g. human keratin, pig trypsin).

**Table 1.** Summary of proteins identified in the saliva of *Heliconius melpomene* via LC-MS/MS.

| Protein | Putative Function | Average Percent Coverage | Unique Spectra | Signal Peptide | Biological Replication | Technical Replication |
| --- | --- | --- | --- | --- | --- | --- |
| **Proteolysis** |  |  |  |  |  |  |
| HMEL003607-PA | trypsin-like protease | 4.925 | 2 | N | 1 | 2 |
| HMEL006217-PA | trypsin-like protease | 22.5 | 19 | U | 2 | 4 |
| HMEL007847-PA | carboxypeptidase | 6.55 | 5 | N | 2 | 4 |
| HMEL010316-PA | astacin | 9.9 | 6 | Y | 2 | 4 |
| HMEL015078-PA | astacin | 9.8 | 7 | Y | 2 | 4 |
| HMEL013718-PA | trypsin inhibitor | 18.75 | 3 | Y | 2 | 3 |
| HMEL017107-PA | trypsin-like protease | 15 | 5 | N | 2 | 4 |
| HMEL002374-PA | cysteine protease inhibitor | 22 | 3 | Y | 2 | 4 |
| HMEL007577-PA | cysteine protease inhibitor | 32.25 | 4 | Y | 2 | 4 |
| HMEL014517-PA | cysteine protease | 22.5 | 5 | Y | 2 | 4 |
| **Carbohydrate hydrolysis** |  |  |  |  |  |  |
| HMEL005611-PA | β-fructofuranosidase | 23.75 | 16 | Y | 2 | 4 |
| HMEL005612-PA | β-fructofuranosidase | 10 | 9 | Y | 2 | 4 |
| HMEL011728-PA | glycerophosphoryl diester phosphodiesterase | 0.625 | 2 | Y | 1 | 1 |
| HMEL014479-PA | hydrolase | 6.5 | 9 | U | 2 | 4 |
| HMEL014593-PA | β-hexosaminidase | 1.55 | 2 | N | 1 | 1 |
| **Immunity** |  |  |  |  |  |  |
| HMEL005769-PA | REPAT gene | 39.5 | 5 | Y | 2 | 4 |
| HMEL013458-PA | hemolin | 1.2 | 2 | Y | 1 | 1 |
| HMEL010053-PA | GMC oxidoreductase | 2.9 | 3 | U | 2 | 4 |
| HMEL010482-PA | alpha crystalline / HSP20 | 31.5 | 8 | Y | 2 | 4 |
| HMEL002661-PA | lysozyme | 21 | 4 | Y | 2 | 4 |
| HMEL016918-PA | $\beta$ 1,3 glucanase | 2.325 | 2 | Y | 1 | 2 |
| **Housekeeping or other function** |  |  |  |  |  |  |
| HMEL002092-PA | yellow-d | 3.55 | 2 | Y | 1 | 2 |
| HMEL010248-PA | Splicing factor | 1.53 | 2 | N | 2 | 2 |
| HMEL013620-PA | actin | 5.2 | 3 | N | 2 | 4 |
| HMEL015393-PA | CAP domain | 37 | 3 | U | 2 | 4 |
| **Unknown Function** |  |  |  |  |  |  |
| HMEL008913-PA | Unknown | 38.5 | 4 | Y | 2 | 4 |
| HMEL008915-PA | Unknown | 24 | 4 | Y | 2 | 4 |
| HMEL014907-PA | Unknown | 7.5 | 2 | Y | 1 | 2 |
| HMEL015039-PA | Unknown | 17 | 3 | Y | 2 | 4 |
| HMEL015041-PA | Unknown | 15 | 2 | Y | 2 | 4 |
| HMEL010245-PA | Unknown | 7.025 | 3 | Y | 2 | 4 |

One clear prediction about salivary proteins is that they are secreted extracellularly and therefore should contain a signal peptide at the N-terminus (Scheele et al., 1978). As expected, signal peptides predicted by Signal-P (via InterproScan) were found in 20 of the salivary proteins (Petersen et al., 2011). This is probably an underestimate because four of 11 proteins without predicted signal peptides were represented by problematic gene models that lacked start codons. Missing start codons likely reflects errors in the underlying genome assembly on which gene models were built because manual inspection could not identify obvious start codons. Otherwise, “complete” proteins without signal peptides tended to have “housekeeping” functions and are likely to be *Heliconius*-derived contaminants rather than true salivary proteins (see section below on *“housekeeping”* proteins).

The identified proteins could be divided into four groups based on function: proteolysis, carbohydrate hydrolysis, immunity, and “housekeeping”. Additionally, several proteins could not be functionally annotated and were lumped into a fifth group of proteins with unknown function.

### 3.3 Proteolytic proteins

Ten identified proteins were found to play a role in proteolysis, encompassing a range of functions including digestion of whole amino acids, cleaving small peptide bonds, and proteolytic inhibition. The seven proteases are primary candidates for playing a role in the digestion of pollen granules. These include serine proteases, cysteine proteases, astacins, and a carboxypeptidase. All three serine proteases appear to have trypsin-like or chymotrypsin-like properties based on BLAST-based homology and protein domain predictions. Intriguingly, both HMEL006217-PA and HMEL017107-PA show close homology (i.e. strong BLAST hits) to the Cocoonase protein from *Bombyx mori* (silkworm). Cocoonase is a well-characterized trypsin-like protease secreted by the proboscis during eclosion to weaken the cocoon silk and facilitate emergence (Kafatos et al., 1967; Yamamoto et al., 1999). The function of Cocoonase homologs in butterflies, which lack silken cocoons, remains unknown. In the case of *Heliconius* it is tempting to speculate that these proteases, which presumably have an evolutionary history of expression in the proboscis, were evolutionarily co-opted to function in pollen digestion.

Carboxypeptidases hydrolyze peptide bonds at the carboxy-terminal end of a peptide or protein and are also known for their digestive roles (Bown and Gatehouse, 2004). Similarly, astacins often play an important role in extracellular protein digestion (Foradori et al., 2006). Thus this suite of secreted proteases together potentially provides a rich cocktail for breaking down pollen proteins and releasing free amino acids for consumption.

The cysteine and the trypsin inhibitors inactivate the cysteine and serine proteases, respectively, by bonding to the protein’s active site and rendering it inactive (Eguchi, 1993). The two cysteine protease inhibitors identified here appear to be related to the well-characterized Bombyx Cysteine Protein Inhibitor (BCPI) (Yamamoto et al., 1999). BCPI-like proteins likely originated from the inhibitory propeptide region of a cysteine proteinase that is typically cleaved to release the proteolytic function of the mature peptide. These BCPI-like proteins function as “stand alone” inhibitors of cathepsin-L type cysteine proteases (Kurata et al., 2001). Another such protein was proteomically identified as a constituent of seminal fluid in *Heliconius erato*; the putative *H. melpomene* ortholog of this seminal protein is clearly distinct from these two salivary cysteine protease inhibitors, sharing only ~70% amino acid identity with either (Walters and Harrison, 2010). It thus appears that these propeptide-derived cysteine protease inhibitors are commonly deployed for extra-cellular regulation of proteolysis in *Heliconius* butterflies. Nonetheless, it is difficult to predict what role, if any, these cysteine and trypsin protease inhibitors play in pollen digestion.

A distinct lack of molecular characterization of other butterfly saliva proteomes leads to difficulty in making comparisons across pollen and non-pollen feeding Lepidoptera. However, a study performed by (Feng et al., 2013) gave insight into the honeybee saliva proteome. Honeybees are another insect that consumes both pollen and nectar, presenting interesting parallels to *Heliconius*. Honeybees have a mostly carbohydrate rich diet (nectar), which is reflected in the proteins found in their proteome. Both proteomes contain proteins relating to both proteolytic activity and carbohydrate hydrolysis, but *Heliconius* appears to have relatively more proteins related to proteolytic activity and fewer to carbohydrate hydrolysis.

### 3.4 Carbohydrate hydrolysis

Five proteins identified in *H. melpomene* saliva are predicted to be varieties of glycoside hydrolases that appear to play a role in carbohydrate hydrolysis (Withers, 2001). The two β-fructofuranosidases function in breaking down sucrose into fructose and glucose by cleaving the O-C bond. Until recently, β-fructofuranosidases were thought to be absent from animals despite being found among bacteria, fungi, and plants. However, pairs of these proteins have been identified in several lepidopteran species, apparently having arisen via horizontal transfer from bacteria (Daimon et al., 2008). Previously, these β-fructofuranosidases have primarily been associated with larval gut, so their presence in adult saliva is consistent with a role in digestion but also marks a distinct expansion of their known functional milieu.

The remaining three glycoside hydrolases (glycerophosphodiester phosphodiesterase, β-hexosaminidase, and hydrolase) all appear to have relatively general functions in sugar metabolism. This is not unexpected given that *Heliconius* butterflies consume substantial quantities of sugar-rich plant nectar along with pollen.

### 3.5 Immune function

Another six *H. melpomene* salivary proteins likely play a role in immune response. Two of these, lysozyme and β-1, 3 glucanase, are glycoside hydrolases that have secondarily evolved to function in immune response (Davis and Weiser, 2011). Lysozymes are common antimicrobial proteins that function to degrade bacterial cell walls; they are well known components of insect immune responses, including in Lepidoptera (Callewaert and Michiels, 2010; Jiang et al., 2010). Proteins that bind β-1,3glucan function as pathogen recognition proteins that tend to target gram-negative bacteria. Several such proteins have been identified in moths and butterflies (Fabrick et al., 2004). These proteins are usually isolated from hemolymph, but have also been found in the saliva and digestive tracts of other insects (Pauchet et al., 2009).

REPAT and hemolin are Lepidopteran specific immune proteins that have shown increased expression in response to pathogen infection in caterpillars of several species (Hernández-Rodríguez et al., 2009; Terenius et al., 2009; Yamamoto et al., 1999). Also implicated in insect immune response are heat shock proteins, such as alpha crystalline, that are important in keeping essential proteins from unfolding (Pirkkala et al., 2001). Hsp20/alpha crystalline has been found in the salivary glands of other insects and is known to regulate proteins when the organism’s temperature exceeds 25 degrees C (Arrigo and Ahmadzadeh, 1981). Finally, we have tentatively assigned an immunity-related function to the one identified salivary glucose-methanol-choline (GMC) oxidoreductase gene. GMC oxidoreductases comprise a large and diverse protein family whose members play a variety of often poorly understood roles in developmental processes, glucose metabolism, and immune function (Iida et al., 2007). In Lepidoptera this protein family is particularly diverse and many members seem to play a role in immune response (Sun et al., 2012). Thus we have grouped this protein with other immunity-related proteins, but much additional research would be necessary to confidently characterize the true function of this particular GMC oxidoreductase.

### 3.6 *Housekeeping* and other functions

Proteins functioning in proteolysis, sugar metabolism, and immunity are reasonably expected to be found in saliva. We additionally identified in our samples several proteins that seemingly have little relevance to expected salivary functions, or are generally of ambiguous function. Foremost among these is actin, known for its role in muscle contraction and cytoskeletal structure generally, but not expected to function outside of cells (Dominguez and Holmes, 2011). Actin is a ubiquitous and highly abundant protein, so may easily have contaminated the saliva samples. Similarly, an identified serine-arginine-rich splicing factor protein typically functions in RNA splicing and gene expression (Long and Caceres, 2009); it is also probably best considered a contaminant.

Somewhat more ambiguous is the presence of yellow-d, a member of the *yellow* protein family. The function of Yellow proteins is poorly understood, though clearly some members play a role in melanization (Drapeau, 2001; Ferguson et al., 2010). In *B. mori*, yellow-d appears to be ubiquitously expressed and also contains a predicted signal peptide (Xia et al., 2006). The annotation of the yellow-d gene model from the *H. melpomene* genome did not indicate the presence of a signal peptide. However, comparison with a sequence generated from ESTs (GenBank accession ADX87351) clearly indicates that the genome-based model is truncated and that *H. melpomene* yellow-d does contain a signal peptide. Thus, while the molecular function of this and other yellow proteins remains largely unknown, it seems reasonable to consider yellow-d as normally present in *H. melpomene* saliva.

The Cysteine-rich secretory proteins, antigen 5 and pathogenesis related (CAP) proteins are taxonomically diverse with an equally diverse set of functions, making it difficult to predict any particular function for this one protein found in H. melpomene saliva (Gibbs et al., 2008). CAP proteins are typically secreted extracellularly, but in the case of this one salivary CAP, the predicted gene model was incomplete at the N-terminus and therefore uninformative regarding the presence of a signal peptide.

### 3.7 Unknown function

Finally, six proteins found in the sample could not be functionally characterized at any level, other than all of them exhibiting a predicted signal peptide. One of these, HMEL010245-PA, showed extensive homology to similar proteins present in many other insect species, though none of these were functionally annotated. The remaining five proteins appear to be extremely taxonomically restricted. HMEL015039-PA and HMEL015041-PA are a pair of closely linked paralogs situated adjacent to each other, separated by ~8Kbp, suggesting they arose via tandem duplication. Strikingly, a variety of BLAST strategies have yielded no significant homology (e-val < 0.01) to any other protein or nucleotide sequences.

The remaining three uncharacterized proteins, HMEL008913-PA, HMEL008915- PA, and HMEL014907-PA, are another set of paralogs. The similarity and apparent tandem duplication of HMEL008913-PA and HMEL008915-PA suggest HMEL014907-PA is the most distantly related of the three paralogs. In this case, the only clearly homologous loci that were identified were a pair of paralogs from the monarch butterfly, KGM_02914 & KGM_02913, that also appear to be tandemly duplicated. Otherwise these proteins lacked both Blast and InterproScan hits, although each had a signal peptide. These groups of Nymphalid-specific, perhaps even *Heliconius*-specific, secreted proteins in the saliva are very intriguing in light of *Heliconius* pollen feeding. Further investigation of the origin and function of these proteins, along side the other better-characterized salivary proteins we have identified, will be essential for comprehensively understanding the evolutionary novelty presented by *Heliconius* pollen feeding.

## 4 Acknowledgements

This work was supported by the Balfour-Browne fund administered by the Department of Zoology at the University of Cambridge. Additional support came from the University of Kansas. We are grateful for technical assistance from Nadya Galeva and Todd Williams from the University of Kansas proteomics facility. Timothy Karr provided valuable guidance interpreting proteomic data.

